# Beam search decoder for enhancing sequence decoding speed in single-molecule peptide sequencing data

**DOI:** 10.1101/2023.07.13.548796

**Authors:** Javier Kipen, Joakim Jaldén

**Affiliations:** Division of Information Science and Engineering, Kungsliga Tekniska Högskolan, Stockholm, Stockholm, Sweden

## Abstract

Next-generation single-molecule protein sequencing technologies have the potential to accelerate biomedical research significantly. These technologies offer sensitivity and scalability for proteomic analysis. One auspicious method is fluorosequencing, which involves: cutting naturalized proteins into peptides, attaching fluorophores to specific amino acids, and observing variations in light intensity as one amino acid is removed at a time. The original peptide is classified from the sequence of light-intensity reads, and proteins can subsequently be recognized with this information. The amino acid step removal is achieved by attaching the peptides to a wall on the C-terminal and using a process called Edman Degradation to remove an amino acid from the N-Terminal. Even though a framework (Whatprot) has been proposed for the peptide classification task, processing times remain restrictive due to the massively parallel data acquisicion system. In this paper, we propose a new beam search decoder with a novel state formulation that obtains much lower processing times with slightly higher accuracies than Whatprot. Furthermore, we explore how our novel state formulation may lead to even faster decoders in the future.

**Author summary:** Proteomic analyses are often carried on with mass spectrometry, but this method cannot identify low-abundance proteins. Single-molecule protein sequencing methods can overcome this issue, and fluorosequencing is one of these technologies. Fluorosequencing has attracted interest from investors, as evidenced by the recent funding of Erisyon, a company developing this technology. This technique contains a challenging classification task: determining the original peptide sequence from light-intensity observations obtained after several Edman cycles. A classifier based on a combination of *k* Nearest Neighbors (*k*NN) with Hidden Markov Models (HMM) had been shown to have close-to-optimal accuracy with tractable complexity. We propose in this paper a new algorithm that not only improves accuracy compared to state-of-the-art methods but also reduces computation time.

## Introduction

Single-molecule protein sequencing techniques are one of the seven technologies which will have an outsized impact in 2023, according to Nature [1]. These technologies can obtain higher sensitivity on low-abundance proteins [2], which is an advantage compared to mass-spectrometry methods. This utility could enable more sensitive diagnostics [3] and could be a breakthrough in single-cell proteomics. Nowadays, many techniques are being researched and developed [4] [5] [6], and they usually exploit concepts of DNA/RNA sequencing. It is worth mentioning that some of the methods already received capital for industry development [7] [8], and one enterprise has already started delivering the first next-generation single-molecule protein sequencing system [9].

One of the most promising methods is Fluorosequencing [10], which is already in a stage of industry adoption [7]. Flurosequencing can sequence millions of peptides in parallel and quantify them on a zeptomole scale [11]. However, the process still has challenges that make the decoding of the reads a complex task. Among other challenges, the reagents utilized in the Edman degradation [12] [13] affect the chemical attachment of the fluorophores, and the rate of failure of the Edman degradation is considerable.

Smith et al. introduce in [14] a model-based decoder to obtain the most likely peptide given the light intensity reads. First, Smith et al. propose a HMM that considers all the significant failures during the process and has optimal accuracy. However, this method has untractable computing times for a large number of peptides, so Smith et al. present a hybrid model. This hybrid method combines the HMM with a *k* Nearest Neighbours (*k*NN) pre-filter, along with some other optimizations which allow it to run in tractable times with no significant loss in accuracy.

In this paper, we introduce a beam search decoder (probeam) for the same problem, with a new formulation of states. These states group together a considerable amount of peptides for the first reads, but become more specific as the degradation cycles occur. The proposed beam search decoder has a user-set variable N_B_ which is the number of most likely states kept at each cycle. Our decoder achieves much faster computation times and also obtains slightly higher accuracy than the hybrid model of [14]. Although probeam has a poorer estimation of the probability of the picked sequence, it can achieve 100x faster computing times when decoding using a dataset of twenty thousand proteins.

### The problem

#### Fluorosequencing

Fluorosequencing is a method for sequencing peptides at the level of single molecules [11] [10]. This single-molecule protein sensing technology first breaks the protein into short peptides. Secondly, fluorescent molecules which attach to particular amino acids are added, and then the peptides are attached to a surface. Afterward, an amino acid at a time is removed via Edman degradation [12] [13], a chemical process that removes an amino acid from the N-terminal. By observing the changes in the color intensities, it can be obtained which peptide was the original. Finally, one can determine which protein was the original given the peptides found.

Although it is a promising method, several parts of the process have considerable failure rates, which make the peptide decoding task harder. For example, it is possible that some dyes do not attach to the respective amino acids before anchoring (dye miss), the Edman degradation can be unsuccessful (not removing the last amino acid), dyes can detach between each cycle (dye loss), and the whole peptide can detach spontaneously from the wall. In addition, there are a considerable amount of sequences to decode in practice. This situation means that the peptide decoding algorithm should be fast to not be the process bottleneck.

In this paper, only the decoding problem of classifying the dye sequence from the light intensity reads is analyzed. The task of retrieving the protein from the estimated peptides is out of the scope of this paper, but there are several methods of mass spectrometry that can be reutilized [15] [16].

#### Considerations

The number of colors dyes is referred to as N_D_, and the number of cycles of the Edman degradation cycle is named N_C_. A sample read is called ***X*** where 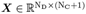. We refer to the observation at step t as 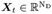 for 0 ≤ *t* ≤ N_C_, so that 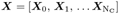

The model parameters are shown in Table 1. The values of the parameters are the same as the ones used in [14]. This paper assumes the same parameters along all colors, as was also done in [14], but the method can be easily extended to color-specific parameters.

**Table 1.**
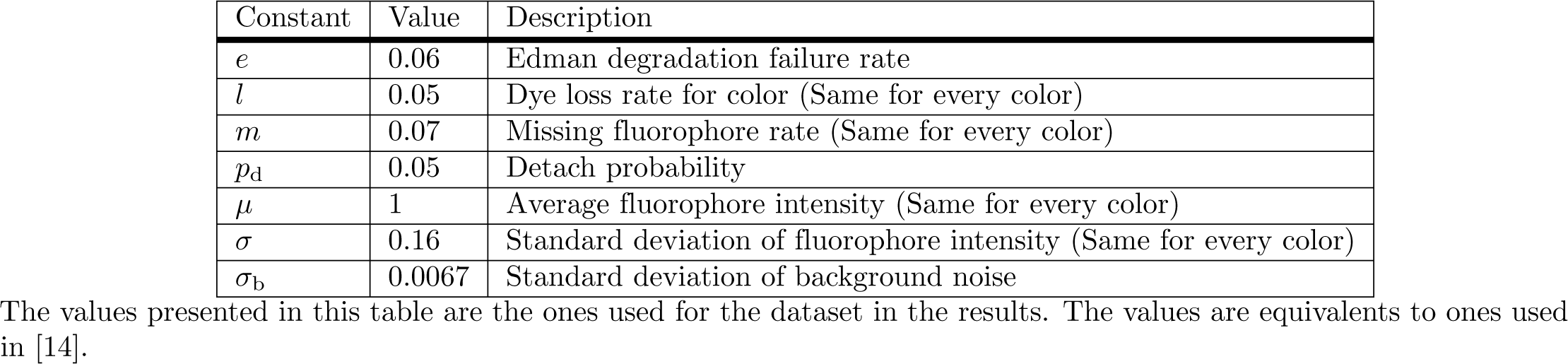
Parameters of the process considered.

| Constant | Value | Description |
| --- | --- | --- |
| $e$ | 0.06 | Edman degradation failure rate |
| $l$ | 0.05 | Dye loss rate for color (Same for every color) |
| $m$ | 0.07 | Missing fluorophore rate (Same for every color) |
| $p_d$ | 0.05 | Detach probability |
| $\mu$ | 1 | Average fluorophore intensity (Same for every color) |
| $\sigma$ | 0.16 | Standard deviation of fluorophore intensity (Same for every color) |
| $\sigma_b$ | 0.0067 | Standard deviation of background noise |

The values presented in this table are the ones used for the dataset in the results. The values are equivalents to ones used in [14].

Every peptide is a sequence of amino acids. Every amino acid can be represented with a letter; then, the peptide can be represented by a sequence of letters. [14] defines the dye sequence representation of a peptide by representing each amino acid by the index of the color that can attach to it. If no dye attaches to that amino acid, a ”.” is assigned. Additionally, in a dye sequence the ”.” after the last index of a color are erased. This removal is because the sequencing past the last dye color would be non-informative. An example of this representation is shown in Table 2.

**Table 2.**
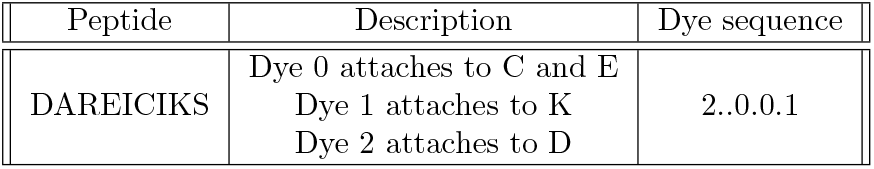
Dye sequence representation example.

With these notations, some peptides can have the same dye sequence and be impossible to distinguish in the measuring system. The decoding task becomes then dye sequence classification instead of peptide classification. The variable ***Y*** represents the true dye sequence that generated the read ***X***. The set of all possible dye sequences is denoted 𝔇_s_, then ***Y*** *∈* 𝔇_s_.

A dye sequence classificator then is a function 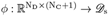. We introduce the random variable *D ∈* 𝔇_s_, representing the randomly selected dye sequence. Then we define the prior distribution over *D* as P_*D*_(*d*). Since the sequences of the proteins and how the protease cuts them is known, P_*D*_(*d*) is known.

## Results

### Dataset

Different datasets were created using the Github code from [14]. First, a dataset of a hundred thousand reads for different amounts of proteins, ranging from twenty proteins to twenty thousand proteins, was generated. This dataset is used to compared the accuracy and running times of the different classifiers. The first time the datasets were generated, only the parameters mentioned in Table 1 were necessary. However, after the commit ”25750c9” of the Whatprot implementation in GitHub [14], two parameters were added that are not mentioned in [14]. These parameter probabilities were set to zero for this paper.

Another dataset was created with one million reads but only from a thousand proteins. This dataset was used to compare the accuracy of the beam decoder against the HMM from [14].

It is also worth noting that the code to generate the dataset was run with the number of reads stated above, but only around 92% of the reads are stored because measurements that start with no dyes attached are removed.

### Performance comparison

Using the first dataset mentioned in the previous section, the accuracy and prediction time of the beam decoder and the state-of-the-art method introduced in [14] as Whatprot were compared in Fig. 1 for a different number of proteins. In the accuracy plot, the dataset was split into ten sub-datasets to estimate the mean and standard deviation of the accuracy estimator. The barely visible error bars in the accuracy plot denote the mentioned standard deviation. Also, the decoding with Whatprot was made with only one thread to compare with an even computational resource. The proposed beam decoder could be easily parallelized across different sample reads, so the correct way to compare them is by running with only one thread.

**Fig 1.**
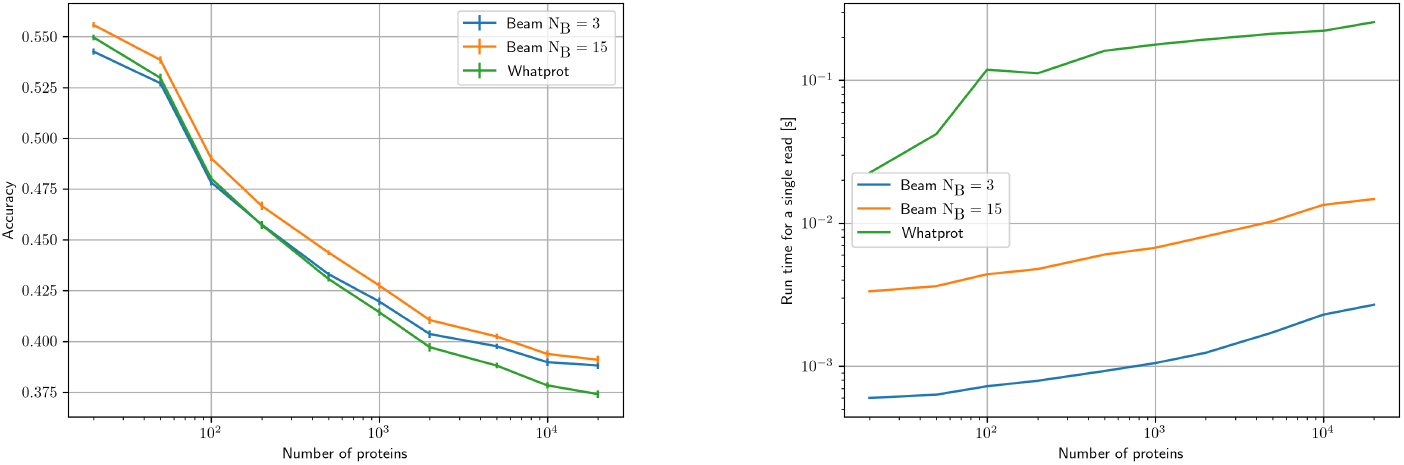
(Left) Accuracy comparison. We plot the accuracy in the classification task for different amounts of proteins and different decoding methods. The bars represent the standard deviation of the estimated accuracy. **(Right) Comparison of run time**. We plot the average prediction time for a single read for both methods.

Whatprot was run with the default parameters used in the results of [14]: 10000 neighbors for the *k*NN, *σ* = 0.5 for the Gaussian weighting function for neighbor voting, a cutoff of *H* = 1000 max peptides from the *k*NN pre-filter and *p* = 5 as the pruning cutoff for HMM.

The beam decoder only has the parameter of the beam width N_B_. After an analysis of the optimal N_B_ for twenty thousand proteins proteins done in Table, the parameter was set to N_B_ = 3 since it had better accuracy and lower runtime. We also included a case with a big number of beams (N_B_ = 15). This number of beams was selected since the accuracy does not increase significantly in the twenty thousand proteins dataset when increasing the number of beams.

It can be observed in Fig 1 that the Beam decoder has a much lower prediction time and a slight improvement in classification accuracy at the same time. If we compare at twenty thousand proteins proteins, we observe approximately a 100x improvement in computing time. This result shows that this classifier is more efficient than the state-of-the-art method. Fig 1 also shows that with a higher number of beams, better accuracy can be obtained, and it is still considerably faster than Whatprot.

### Probability estimates from the beam decoder

We run the HMM classifier from [14] without any pruning and the probeam for the dataset of one thousand proteins to compare the probability estimates. Then the reads which were classified correctly by both the HMM classifier and probeam were extracted. The probability estimations of the dye sequence were kept for each method. Fig 2 shows a 2D-histogram comparing the estimated probability for the different methods on the reads mentioned above.

**Fig 2.**
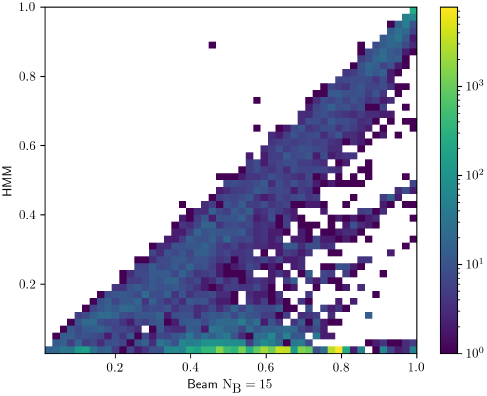
Accuracy estimation. We plot in this 2D-histogram the predicted dye sequence probability of the beam decoder vs the true dye sequence probability, obtained with the HMM from [14]

It can be observed that the estimated probability of the beam decoder is not an accurate measurement of the probability of the picked sequence being correct. Generally, probeam overestimates the score of the chosen peptide because it keeps a smaller amount of possible dye sequences in the very last steps of the decoding. This shows the trade-off when using the proposed beam decoder: faster computing times and higher accuracy can be obtained at the expense of a poorer estimation of the probability. Future work could be to investigate this probability estimation to see if it can be enhanced.

### Beam decoder accuracy

In the second dataset, the beam decoder and the HMM were run to compare the accuracy obtained. As this dataset is bigger, it allows for less variance in estimating our accuracies. The accuracies of the HMM and the beam decoder are shown in Table 3. The HMM was run without pruning, so there were no parameters set, while the beam decoder was run with N_B_ = 15 as in the previous sub-section, since a higher number of beams did not show higher accuracies in the twenty thousand proteins dataset.

**Table 3.**
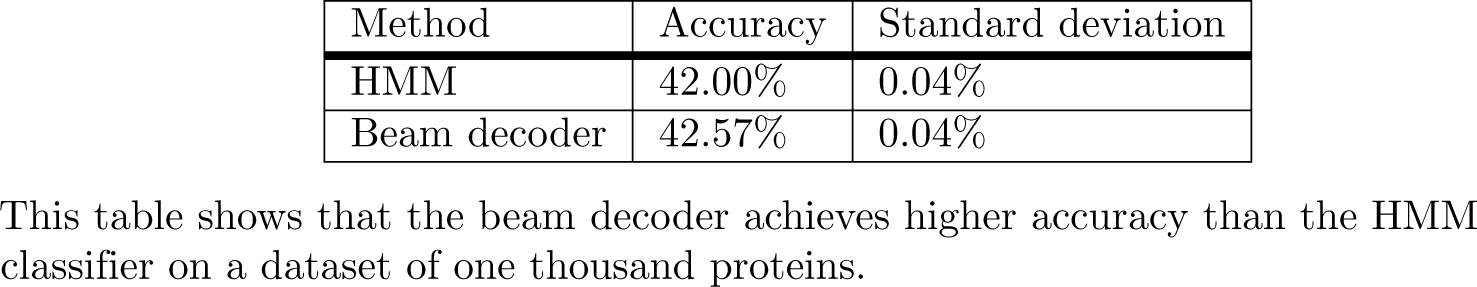
Accuracy comparison to HMM.

This table shows that the beam decoder achieves higher accuracy than the HMM classifier on a dataset of one thousand proteins.

The HMM model without pruning implements the MAP decision rule, which minimizes the error rate. This means that this accuracy should be the highest achievable. However, we can observe that the beam decoder actually has a slightly higher accuracy, and it is very unlikely that is due to the randomness of the simulation since the standard deviation of the accuracy estimator is significantly lower than the accuracy gap. One possible explanation could be that Whatprot is more sensitive to errors in numerical computation than probeam, leading to decreased accuracy.

## Discussion

We propose an algorithm based on the beam search algorithm and a novel state definition for peptide classification from light intensity measurements within fluorosequencing. Our method outperforms state-of-the-art methods in terms of both speed and accuracy.

Our method was not parallelized within the same read because synchronization would be needed whenever the most likely states were kept. However, fluorosequencing relies on sequencing millions of peptides simultaneously, and our method could be applied in parallel for each read. To ensure a fair comparison then, we compared our method to only one thread of the state-of-the-art decoder Whatprot [14] when assuming fixed computing resources. Further improvements could be made by implementing our method for GPU computing, which would significantly reduce the total prediction time.

We did not consider storing all states and transition probabilities a priori. This was because in the development states with more variables were considered. It remains a challenge because of the ample state space, but one could store some of the most likely transitions and states in memory, reducing the computing time.

This paper did not consider the possibility of using deep learning methods for this classification. Some RNNs (Recurrent Neural Networks) have achieved groundbreaking performances in nanopore DNA sequencing [17] [18]. More recently, a hybrid approach between HMM and RNN has obtained state-of-the-art accuracies in that domain [19]. However, these methods still require an enormous amount of data to be trained, and there is not public access yet to experimental data for fluorosequencing experiments.

Another disadvantage of the neural network based approaches is their unreliability in measuring uncertainty. The beam decoder also has an incorrect estimation of the uncertainty. However, the correction of the uncertainty estimation was out of the scope of the paper, and future work could be to analyze these estimations and improve them.

## Methods

### State-of-the-art decoder (Whatprot)

A Hidden-Markov-Model-based classifier was proposed in [14] to solve the above-mentioned classification task. This decoder can be thought of as a derivation from the MAP decision rule where Bayes’ theorem is applied:

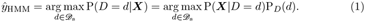

Here P_*D*_(*d*) is known, and P(***X*** | *D* = *d*) can be calculated effectively with a Hidden Markov Models forward algorithm. This method is optimal in terms of accuracy but is computationally expensive. For example, when using the human proteome (≈ twenty thousand proteins proteins), approximately 130 thousand dye sequences need to be considered.

A hybrid method was also introduced in [14]. This method combines the Hidden Markov Model classifier with a prefiltering done with *k* Nearest Neighbours(*k*NN). With *k*NN, the most promising dye sequences are selected (*𝔇*_*k*NN_(***X***)), and then the probabilities of each sequence are obtained again using Bayes’ theorem. This final step assumes that the probabilities of the dye sequences which were not considered are negligible. This Hybrid model classification can be explained then with Equation 2:

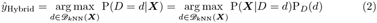

It is worth mentioning that the parameters of the *k*NN were analyzed to have a considerable improvement in processing time without losing significant accuracy in [14]. We refer to this hybrid method when we mention the Whatprot classifier. In addition, the classifier code has other optimizations which make the code run with tractable predicting times:

1. A custom *k*NN is developed, where the reads are discretized, and repetitions are removed and replaced with weights to the distance to the given read.
2. The authors prune the Hidden Markov Models in the forward algorithm with the observation probability. This pruning significantly decreases the processing time at the expense of a minimal drop in accuracy.
3. The transition probabilities are factorized, which results in a much faster way to calculate the forward algorithm
4. The code is written in the C programming language, optimized with data locality and multithreading.

### Beam decoder

#### Inspiration

As we have already written above, Whatprot [14] shows an improvement in computing time without significant losses in accuracy compared to the optimal HMM decoder. However, the calculation time for decoding can be improved.

Our model draws inspiration from the first HMM decoders used for nanopore DNA sequencing [18]. In these models, each state represents a *k*-mer going through the pore, and the observation probabilities depend on this *k*-mer. For the fluorosequencing problem, the observations depend only amount of fluorophores of each color attached in the whole chain ***K***. We therefore included the count of attached fluorophores in each state.

Unlike in the HMM for nanopore base-calling, where transition probabilities were limited to one shift in the *k*-mer and a new base or the same base as before, the fluorosequencing problem’s transition probabilities are more complex. Since we know the possible sequences a priori, we must consider them in the decoding task. We found that we can efficiently calculate these transition probabilities by including two other variables in the states and the a priori information. The first variable, ***N***, represents the number of ideal fluorophores attached. In other words, it is the number of fluorophores the chain would have if *m* = *l* = 0. The second variable *R* is the removed sequence up to that cycle. With these two variables, we can find which are the known sequences that match them and then calculate the transition probabilities.

In this approach, the observation probabilities and the transition probabilities one step ahead are easy to calculate. However, the state space is large. We then can obtain the most likely states via a beam search.

#### State definition

We define the states of our model with three variables shown in Table 4.

**Table 4.** State variables.

| Variable | Description | Example ($N_D = 3$) |
| --- | --- | --- |
| $\mathbf{N}$ | Number of remaining amino acids that attach to each color. $\mathbf{N} \in \mathbb{N}^{N_D}$ | [3 0 2] |
| $\mathbf{K}$ | Number of remaining attached dyes for each color $\mathbf{K} \in \mathbb{N}^{N_D}$ | [1 0 1] |
| $R$ | Removed dye sequence | ".." |

These are the variables that discriminate each state.

We refer with *i* to the *i*th color (0 ≤ *i <* N_D_, *i ∈ ℕ*). Then we define *K*_*i*_ and *N*_*i*_ to the *i*th component of the respective vector (***K*** and ***N***). It is useful to note that:

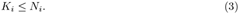

For notation and computation, we consider that if it is detached, then *K*_*i*_ = 0 ∀*i*. Examples of states are shown in Fig. 3.

**Fig 3.**
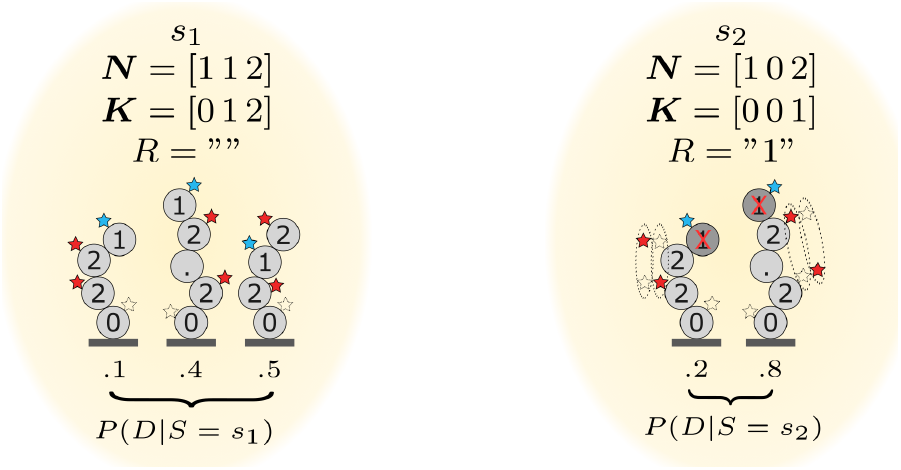
Example of states. *s*_1_ is an example of a state without any removal, and the plotted dye sequences represent all the possible dye sequences for the state in this example. The dotted star means that there is no dye attached. *s*_2_ is another state where there is a removal, and there are different combinations of attached dyes for the color red. One can interpret that if the system evolved from *s*_1_ to *s*_2_ in one degradation cycle, it means that the degradation worked successfully, a ”1” was removed and a ”2” was lost

The states in Fig. 3 are represented only by the variables ***N***, ***K***, and *R*. The paper uses the notation *s*.***N*** as the variable ***N*** of the state *s*, as in objected-oriented programming. We first introduce some functions and definitions to explain how we obtain the corresponding dye sequences for each state.

First, we define the dye-counting function 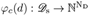 as a function that counts how many times each color appears on a dye sequence. One example is *φ*_*c*_(”1.02..0.0.2”) = [3, 1, 2], and the formal definition is

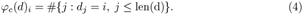

We then define a function *f* (*r, d*) that indicates whether the dye sequence *d* starts with a given sequence *r* or not :

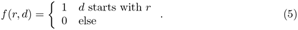

With the previous functions, we can now define the function *η*(***N***, *R*) in Equation 6. This function shows if a dye sequence belongs to one state. Note that it only depends on the variables ***N*** and *R*. In the case of an empty *R* is easy to interpret, the corresponding dye sequences of a state are only the ones whose count is the same as ***N***. When *R* is not empty, one must also check that the dyes started with the same removed sequence.

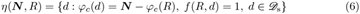

Finally, the relative probabilities of the dyes can be obtained with the information from the a priori distribution:

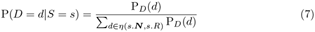

#### Algorithm description

The algorithm can be derived from the MAP decision rule, as Whatprot [14]. In this case, we can rewrite the MAP decision rule by marginalizing a joint distribution of the dye sequences and the states at the last observation 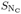, conditioned on the observations

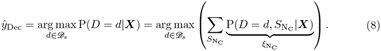

The distribution 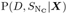 is hard to compute, but we can obtain it recursively. We define *ξ*_*t*_(*d, s*) = P(*D* = *d, S*_*t*_ = *s*|***X***_0_, ***X***_1_ … ***X***_*t*_) so

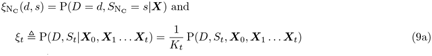

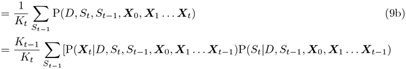

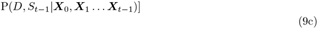

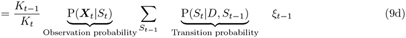

Where 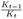 is a normalization factor. First Bayes’ theorem was applied in Equation 9a, and then the previous state was added to the joint, and then the joint was marginalized with respect to *S*_*t−*1_ in Equation 9b. It is important to remark that to obtain Equation 9d the observation probability is assumed to depend only on the state. Nonetheless the transition probability depends on the distribution of the dye sequences too. The Equation 9d is similar to the recursion in the HMM forward algorithm, but it depends on *D*. The observation and transition probabilities are explained in the following sections.

Fig. 4 shows how the random variables are related in our model. This graph indicates that each state does not depend only on other states like in an HMM but also depends on the randomly selected dye sequence, because of the transition probabilities. However, the observations depend only on the state.

**Fig 4.**
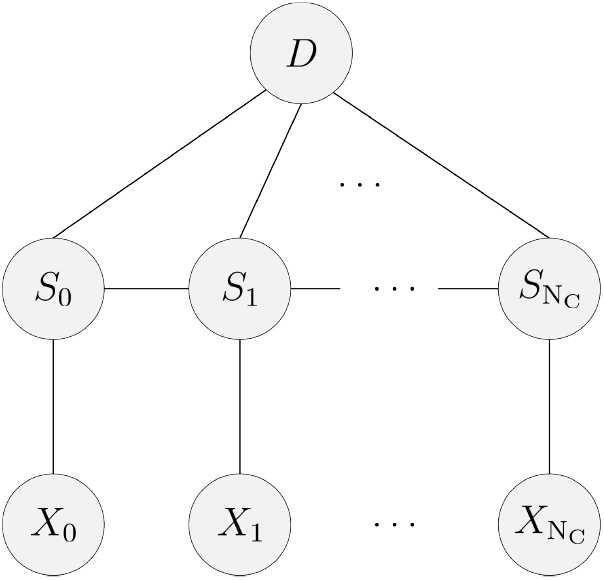
Graphical model. This graphical model represents the dependencies between the states, observations and the randomly picked dye sequence distribution in our model.

An initial step is needed to start the recursion:

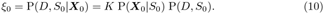

Here the observation probabilities are calculated as in the recursion step. On the other hand, P(*S*_0_) is obtained from a deterministic function applied to P_*D*_(*d*).

This method can obtain the optimal MAP argument, but the number of states considered makes it impossible to calculate. Therefore, this algorithm is implemented keeping only the most likely states at each time step Ŝ_*t*_. After obtaining the initial probabilities with 10, the user-defined number of most likely states N_B_ indicates how many of the most likely states are kept. In the recursion also the N_B_ most likely states are kept.

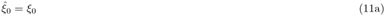

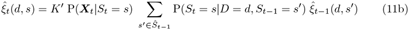

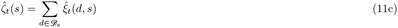

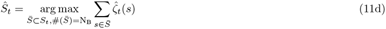

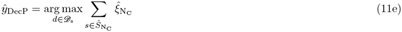

First, the joint distribution with the first observations *ξ*_0_ is obtained. Then this is marginalized in Equation 11c with respect to the dye sequences to obtain the probability of the initial states. The step Equation 11d picks the N_B_ most likely states, and the recursion continues with Equation 11b. In Equation 11b the joint is calculated only considering the previous most likely states. The steps of marginalizing the dye sequences to obtain the state probabilities and keeping the most likely are also done in each degradation step. Finally, the most likely dye sequence is picked with Equation 11e.

It is worth noticing that the value of N_B_ can be tuned. There is a trade-off between computing time and accuracy: when N_B_ lowers, computing time is lowered at the expense of a decrease in accuracy. If N_B_ is large enough, this method is equivalent to the MAP estimator.

#### Calculation of initial states

In order to start the recursion, the joint distributions of the initial states and the dye sequences P(*D, S*_0_) must be calculated. First, the ideal initial states distribution 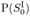 is calculated, which does not consider dye-missing effects. Then the previous results are used to obtain the distribution of the initial states P(*S*_0_). At last, P(*D, S*_0_) is obtained using the previous results.

The ideal initial states are represented as 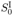. Using the dye counting function (Equation 4) we define the set of possible dye counts 𝔇_N_ as

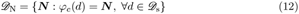

We also define a mapping 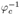 which returns the set of dye sequences whose count is 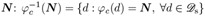. The initial ideal states can be seen as a grouping of the dye sequences that have the same ***N***, so 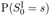 is just the sum of the dye sequence probabilities:

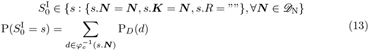

The next step is to consider the missing dye effects, which modify the number of remaining attached fluorophores ***K***. We introduce the set *𝔇*_K_(***N***) which contains all the ***K*** vectors that can be obtained from an initial vector ***N*** because of dye miss or dye loss. These effects can remove any number of fluorophores:

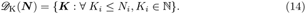

Each configuration of ***K*** due to a dye miss has an associated probability. Since the probability of dye miss is *m* and independent for each fluorophore, P_DyeMiss_(***K*** | ***N***) can be expressed as a combinatorial expression for each color, as was also stated in [14]:

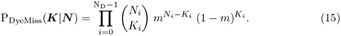

The initial state distribution considers all the possible dye misses for each ideal state. Also, the probability of each initial state can be expressed as the probability of the dye miss times the probability of the ideal state, since both steps are independent:

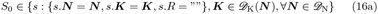

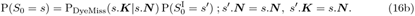

Finally, the joint probability P(*D, S*_0_) can be obtained by applying the conditional probability formula: P(*D, S*_0_) = P(*D*|*S*_0_) P(*S*_0_). P(*S*_0_) was described in the previous steps, and P(*D*|*S*_0_) is defined in Equation 7. Combining these results we obtain:

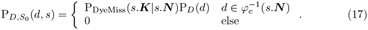

#### Observation probabilities

Then for a state *S*_*t*_ = *s* with a given ***K***, the total standard deviation of fluorophore intensity for the *i*th color 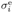 is defined as:

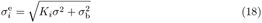

We define the random variable representing the state at time *t* as *S*_*t*_. Using this result, we can easily express the observation probability

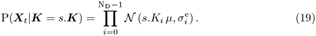

where 𝒩(*a, b*) is a normal distribution with mean *a* and standard deviation *b*. This observation probability was already defined in [14], but here we rewrite it with our notation and our state space.

#### Greedy search for initial states

Equation 17 enables the calculation of the probability of all initial states by marginalizing over the dye sequences, as indicated by Equation 16b. We can express this equation in a more convenient form as

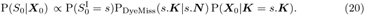

The first two probabilities on the right hand side of Equation 20 can be precomputed offline to optimize runtime, while the third probability depends on the observation. Since we aim to select the most likely states, we can calculate the right hand side of Equation 20 and pick the states which maximize it. One way to solve the initial step is to calculate this probability for every possible initial state. However, this takes considerable computing time and can be done more efficiently without sacrificing optimality.

The observation probability function P(***X***_0_|***K***) defined in Equation 19 is a unimodal function for *K*_*i*_ *∈* ℕ and a fixed ***X***_0_. One can show this by taking the derivative of the expression of Equation 19 with respect to *K*_*i*_ and setting it to zero, and then the only positive solution is the critical point of interest. Evaluating the function’s second derivative with respect to *K*_*i*_ results in a negative expression, indicating that it is a maximum. The critical point is the value of the optimal 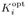, which is a real number. The expressions themselves were not included in this paper due to their cumbersome nature.

Consequently, there exists a value ***K***^opt^ with 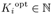 such that P(***X***_*t*_ | ***K***^opt^) ≥ P(***X***_*t*_ ***K***) for all possible ***K***. When considering a state, its fluorophore count ***K*** is bounded by the number of amino acids that attach to each color ***N*** (Equation 3), so ***K***^opt^ may be outside of the bounds. In such cases, we refer to the optimal fluorophore count of a state as 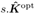, which maximizes the observation probability for a given state, taking into account its bounds.

We initiate the search by creating a list in which we will insert the most likely N_B_ states. First, we initialize the list, and then a search is performed. If a state is found during the search that is more likely than the ones in the list, it is inserted in the list and the less likely state of the list is dropped. To initialize the list, we order the states with the distance from the normalized observations 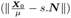. Subsequently, we initialize the list with the N_B_ closest states in this distance, and we obtain their probabilities with Equation 20 evaluating with the optimal fluorophore count of the state 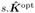.

To facilitate the search process, we introduce the gap variable **∆** to represent the deviation between the fluorophore count *s*.***K*** of a state, and the optimal fluorophore count of it 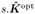 We then perform a breadth-first search for each state where each node represents a deviation **∆**. The multiplication 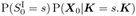 is evaluated in each node with the resulting *s*.***K***. If the mentioned probability is lower than the lowest probability on the list of picked states, it is unnecessary to consider the node or any nodes derived from it. This is because these likelihoods will be equal or lower due to the unimodal property of P(***X***_*t*_ | ***K*** = *s*.***K***). The next step is to calculate the expression in Equation 20 for the nodes that need to be considered. If this value is higher than any of the states in the list, we include this state in the list of N_B_ most likely states, and we drop the less likely.

Fig. 5 illustrates an example of this search process. We begin by testing the optimal ***K***, denoted as 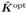, where **∆** = [0, 0, 0]. Subsequently, we evaluate the expression 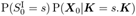 in every node of the same depth. If it is higher than the probability of the less likely state from the list, the node and its neighbors are considered. The breadth-first search allows us to find repeated neighbor nodes and calculate them only once. The mentioned process is repeated until there are no more neighbors or nodes to be considered. It is also worth noticing that next-level neighbors which are also neighbors from discarded nodes (grey states in Fig. 5) can also be discarded. This last optimization was not implemented in our code.

**Fig 5.**
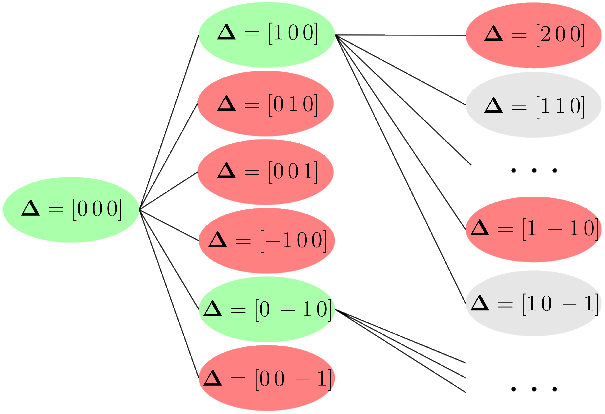
Breadth-first search for initial states. Each node represents the difference **∆** between the optimal fluorophore count and how many fluorophores were attached. On every node, it is evaluated if the neighbors could be candidates or not, and then the nodes are evaluated individually with Equation 20

#### Transition probabilities

We introduce two new variables to show the calculations of the transition probabilities. These meta-states allow us to separate the transition probabilities into three independent phenomena: Edman degradation, dye loss, and detachment. The latter two do not depend on the prior information so that the transition probabilities can be rewritten as:

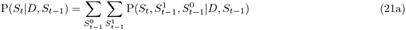

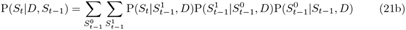

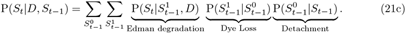

Each phenomenon will be covered separately. It is worth mentioning that these cases are considered only for states that are not already detached or with null ***K***. These states can only transition to themselves with probability 1.

#### Detachment

We define the detach function Det as

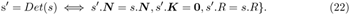

This function removes all the remaining attached fluorophores *s*.***K*** from the input state *s*. Then the probability 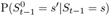 is given by

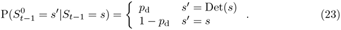

#### Dye Loss

Similar to the equation of dye missing (Equation 15), P_DyeLoss_(***K***^***′***^|***K***) indicates which is the probability of having kept ***K***^*′*^ attached fluorophores from the original ***K* only** due to dye loss within a degradation cycle. Since each fluorophore has an independent probability of dye loss, the probability is a combinatorics formula which has already been stated in [14]:

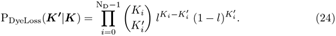

Then we denote with Λ(*s*) as all the possible states that can be obtained only through dye loss from the base state *s*:

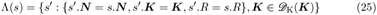

Finally, we can write 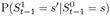 considering that it should be 1 when detached because the dye loss phenomena would not modify the output, and the dye loss probability in the other case:

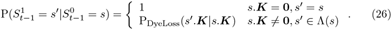

In the practical implementation, we only consider 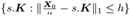 where *h* is a threshold distance set by the user. This step is done to avoid considering very unlikely states. We tested that with *h* = 5 there was no loss in accuracy in a reduced dataset, so it was used. In future implementations, this step could be replaced by a greedy search like in the initial states. This could reduce the number of states considered without losing optimality.

#### Edman degradation

Several functions are described to simplify the expression of probability associated with the Edman degradation.

First, we define functions that decrease *s*.***K*** and *s*.***N*** : Dec_K_ and Dec_N_ respectively. They are used for the possible states after removing a fluorophore of color *i*.

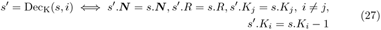

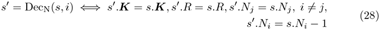

Then the function ℛ(*s, a*) adds the removed amino acid *a* (in dye sequence format) to the removed sequence

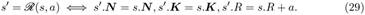

Next, we define the function *α*(*r, d, a*) where R is the removed sequence, *d* is a given dye sequence and *a* the amino acid to remove

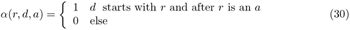

This function states whether removing an amino acid *a* from a sequence *d* that starts with *r* or not is possible. *a* can take the values of the indexes of colors or the value of a dot. Then the probability P_Rem_(*s, a, d*) indicates how likely is to remove an amino acid *a* given the state *s* and the dye sequence with its relative probability:

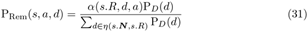

Finally, it is necessary to consider whether the removed amino acid was attached to a fluorophore when a luminescent amino acid is removed. Considering that we have *K*_*i*_ and *N*_*i*_ from the state, there will be 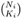 ways to accommodate the dyes to the respective luminescent amino acids. Then we can think of the probability of having the dye in the first luminescent amino acids of all the combinations that pick the first possible a(mino)acid. In order to count them, we will have to pick *K*_*i*_ *−* 1 in *N*_*i*_ *−* 1. This gives us 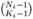 combinations. Therefore the probability of having the Dye Atached in the amino acid to remove P_DA_(*s*) is given by

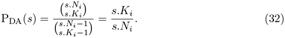

Each case’s sets of possible states will also be identified with different Ψ(*s*). Ψ_n_(*s*) is the state where a dot was removed

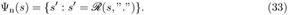

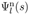 is the set of states where a luminescent amino acid was removed, but the fluorophore was not attached to it

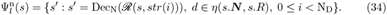

The last possible set is 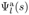, which contains the states where a luminescent amino acid was removed, and the fluorophore was attached to it

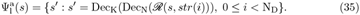

Then the transition probabilities due to Edman degradation are described as

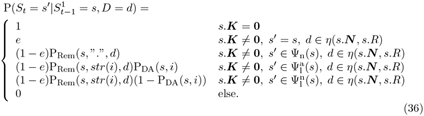

#### Conceptual explanation

Our decoder keeps the most likely states, with each state representing several dye sequences. As the Edman cycles progress, the state space expands, but simultaneously, the states become more specific regarding the represented dye sequences. This feature lets us consider a large number of sequences initially and subsequently narrow down to a discriminative state space.

The states in our model reflect the physical processes occurring during the degradation cycles. At each time step, the possible next states account for peptide detachment, all the combinations of dye loss, and whether the Edman degradation was successful. If degradation is successful, the model considers which dye color was removed and whether a fluorophore was attached, as well as the probability of removing an amino acid that cant attach to a dye.

Although the effects considered in each step lead to a significant increase in the number of possible states, the amount of relevant states remains small. This feature is because the states with a fluorophore count *s*.***K*** at time *t* that deviate greatly from the optimal fluorophore count 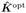 have a significantly lower probability. The unlikelihood arises due to the relatively small fluorophore light intensity’s standard deviation *σ* compared to the fluorophore light intensity’s mean *µ*. Consequently, a small number of states need to be kept in the beam search to achieve near-optimal accuracy, enabling fast computation times.

Whatprot was optimized by minimizing the joint computational time of the pre-filter to select the most likely sequences and the computation of the likelihood of each observation given the candidate sequence. However, their *k*NN pre-filter requires a large dataset to be trained, which results in a considerable computation time to find the candidate sequences. Additionally, even with pruning, the forward algorithm has to consider many states and transitions at a time. This optimized joint computing time is much larger than our model’s computation time. In our model, the recursive step is the most computationally intensive, as it determines the possible next states along with their probabilities and retains the best ones. Furthermore, our method offers potential for further improvement. Among other optimizations that can be done, pre-computing transition probabilities partially during setup time would significantly reduce computing times.

## Supporting information

**S1 Table. Comparison of** N_B_. The values of accuracy and runtime for different values of N_B_ on twenty thousand proteins proteins dataset.

## Acknowledgments

The authors gratefully acknowledge Alicia Cevedo for her proofreading and grammatical corrections to the paper. This work has been supported by the Swedish Research Council (VR Research Environment Grant 2018-06169; QuantumSense), and the Swedish Foundation for Strategic Research (SSF Grant ITM17-0049).

## References

1. Eisenstein M. Seven technologies to watch in 2023; 2023. Available from: https://www.nature.com/articles/d41586-023-00178-y.

2. Restrepo-Pérez L, Joo C, Dekker C. Paving the way to single-molecule protein sequencing. Nature Nanotechnology. 2018;13:786–796. doi:10.1038/s41565-018-0236-6.

3. Kustatscher G, Collins T, Gingras AC, Guo T, Hermjakob H, Ideker T, et al. Understudied proteins: opportunities and challenges for functional proteomics. Nature Methods. 2022;19:774–779. doi:10.1038/s41592-022-01454-x.

4. Alfaro JA, Bohländer P, Dai M, Filius M, Howard CJ, Van Kooten XF, et al. The emerging landscape of single-molecule protein sequencing technologies. Nature methods. 2021;18(6):604–617.

5. Vistain LF, Tay S. Single-Cell Proteomics. Trends in Biochemical Sciences. 2021;46:661–672. doi:10.1016/j.tibs.2021.01.013.

6. Callahan N, Tullman J, Kelman Z, Marino J. Strategies for Development of a Next-Generation Protein Sequencing Platform. Trends in Biochemical Sciences. 2020;45:76–89. doi:10.1016/j.tibs.2019.09.005.

7. Field R. Research bridge partners makes seed capital investment in Erisyon, inc..; 2022. Available from: https://www.researchbridgepartners.org/research-bridge-partners-makes-seed-capital-investment-in-erisyon-in

8. Novet J. Amazon invested millions in a pre-revenue company with a system for measuring human proteins; 2021. Available from: https://www.cnbc.com/2021/08/05/amazon-invested-millions-in-nautilus-biotechnology.html.

9. Quantum-Si’Incorporated. Quantum-Si Reports Fourth Quarter and Fiscal Year 2022 Financial Results; 2023. Available from: https://ir.quantum-si.com/news-releases/news-release-details/quantum-si-reports-fourth-quarter-and-fiscal-year-2022-financial.

10. Swaminathan J, Boulgakov AA, Marcotte EM. A Theoretical Justification for Single Molecule Peptide Sequencing. PLoS Computational Biology. 2015;11. doi:10.1371/journal.pcbi.1004080.

11. Swaminathan J, Boulgakov AA, Hernandez ET, Bardo AM, Bachman JL, Marotta J, et al. Highly parallel single-molecule identification of proteins in zeptomole-scale mixtures. Nature Biotechnology. 2018;36:1076–1091. doi:10.1038/nbt.4278.

12. Edman P, Högfeldt E, Sillén LG, Kinell PO. Method for determination of the amino acid sequence in peptides. Acta chem scand. 1950;4(7):283–293.

13. Edman P, Begg G. A protein sequenator. European Journal of Biochemistry. 1967; p. 80–91.

14. Smith MB, Simpson ZB, Marcotte EM. Amino acid sequence assignment from single molecule peptide sequencing data using a two-stage classifier. bioRxiv. 2023;doi:10.1101/2022.09.23.509260.

15. Käll L, Canterbury JD, Weston J, Noble WS, MacCoss MJ. Semi-supervised learning for peptide identification from shotgun proteomics datasets. Nature Methods. 2007;4:923–925. doi:10.1038/nmeth1113.

16. Keller A, Nesvizhskii AI, Kolker E, Aebersold R. Empirical statistical model to estimate the accuracy of peptide identifications made by MS/MS and database search. Analytical Chemistry. 2002;74:5383–5392. doi:10.1021/ac025747h.

17. ONTplc. Nanoporetech/bonito: A pytorch basecaller for oxford nanopore reads.; 2020. Available from: https://github.com/nanoporetech/bonito.

18. Rang FJ, Kloosterman WP, de Ridder J. From squiggle to basepair: Computational approaches for improving nanopore sequencing read accuracy. Genome Biology. 2018;19. doi:10.1186/s13059-018-1462-9.

19. Xu X, Bhalla N, Ståhl P, Jaldén J. Lokatt: A hybrid DNA nanopore basecaller with an explicit duration hidden Markov model and a residual LSTM network. bioRxiv. 2022;doi:10.1101/2022.07.13.499873.

